# Physiological predictors of competitive performance in CrossFit^®^ athletes

**DOI:** 10.1101/2019.12.16.877928

**Authors:** Rafael Martínez-Gómez, Pedro L. Valenzuela, Lidia B. Alejo, Jaime Gil-Cabrera, Almudena Montalvo-Pérez, Eduardo Talavera, Alejandro Lucia, Susana Moral-González, David Barranco-Gil

## Abstract

The aim of this study was to determine which physiological variables could predict performance during a CrossFit competition. Fifteen male CrossFit athletes (35 ± 9 years) participated and performed a series of tests (incremental load test for full squat and bench press, jump tests, incremental running test, and Wingate test) that were used as potential predictors of CrossFit performance. Thereafter, they performed the five Workouts of the Day (WODs) corresponding to the CrossFit Games Open 2019, and the relationship between each variable and CrossFit performance was analyzed. Overall Crossfit performance (i.e., final ranking considering all WODs) was significantly related to jump ability, mean and peak power output during the Wingate test, relative maximum strength for the full squat and the bench press, and maximum oxygen uptake and maximum speed during an incremental running test (all p<0.05, r=0.58–0.75), although the relationship of most markers varied depending on the analyzed WOD. Multiple linear regression analysis showed that the combination of maximum oxygen uptake, squat jump ability, and reactive strength index accounted for 81% of the variance in overall CrossFit performance (p=0.0003). CrossFit performance seems dependent on a variety of power-, strength-, and aerobic-related markers, which reflects the complexity of this sport. Improvements in aerobic capacity may help people and athletes in CrossFit performance and well-being. Also, focus on lower body power could be the key to obtain better performance markers.

## Introduction

CrossFit is a strength and conditioning exercise program that combines weightlifting, gymnastics, and traditional ‘aerobic’ exercise modalities (e.g., running, rowing, cycling), all of which are performed as quickly as possible within different types of workout sessions – known as “Workout of the Day” (WOD) (1). Despite the popularity of this type of training (1) as well as that of CrossFit competitions (known as Opens, and consisting of five WODs performed consecutively during one month), there is scarce evidence on the determinants of performance in this sport (2–5).

Some studies have found a relationship between markers of maximal aerobic capacity (e.g., maximal oxygen uptake [VO_2max_]) and CrossFit performance (2,4,5). Moreover, we and others (2,3,5) observed that the strongest (e.g., those with the highest 1-repetition maximum [1RM]) and most powerful (e.g., those with the highest power values during the full squat exercise or the Wingate Anaerobic Test [WAnT]) athletes achieved a higher CrossFit performance. However, this relationship seems to be dependent on the type of WOD analyzed (2,4,5).

There is still much controversy about which tests or physiological variables can be used to predict CrossFit performance, and this seems to be partly due to the wide variety of ‘domains’ included in CrossFit WODs (e.g., strength, power and aerobic-related exercises) and the paucity of studies on this topic. Under this context, the aim of the present study was to determine which physiological variables could predict performance during a CrossFit competition (The Open, 2019), by analyzing markers of ‘aerobic’ and ‘anaerobic’ capacity, strength, and power.

## Materials and methods

### Experimental design

During the two weeks prior to the start of the competitive phase (i.e., the CrossFit Games Open 2019), participants performed a series of tests (incremental load test for full squat and bench press, jump tests, incremental maximal running test, and WAnT) to analyze their potential as predictors of CrossFit performance. Thereafter, the participants performed the five WODs corresponding to the CrossFit Games Open 2019 on five different days for one month, always in the same order and on the same day for all participants (Fig 1).

**Fig 1.**
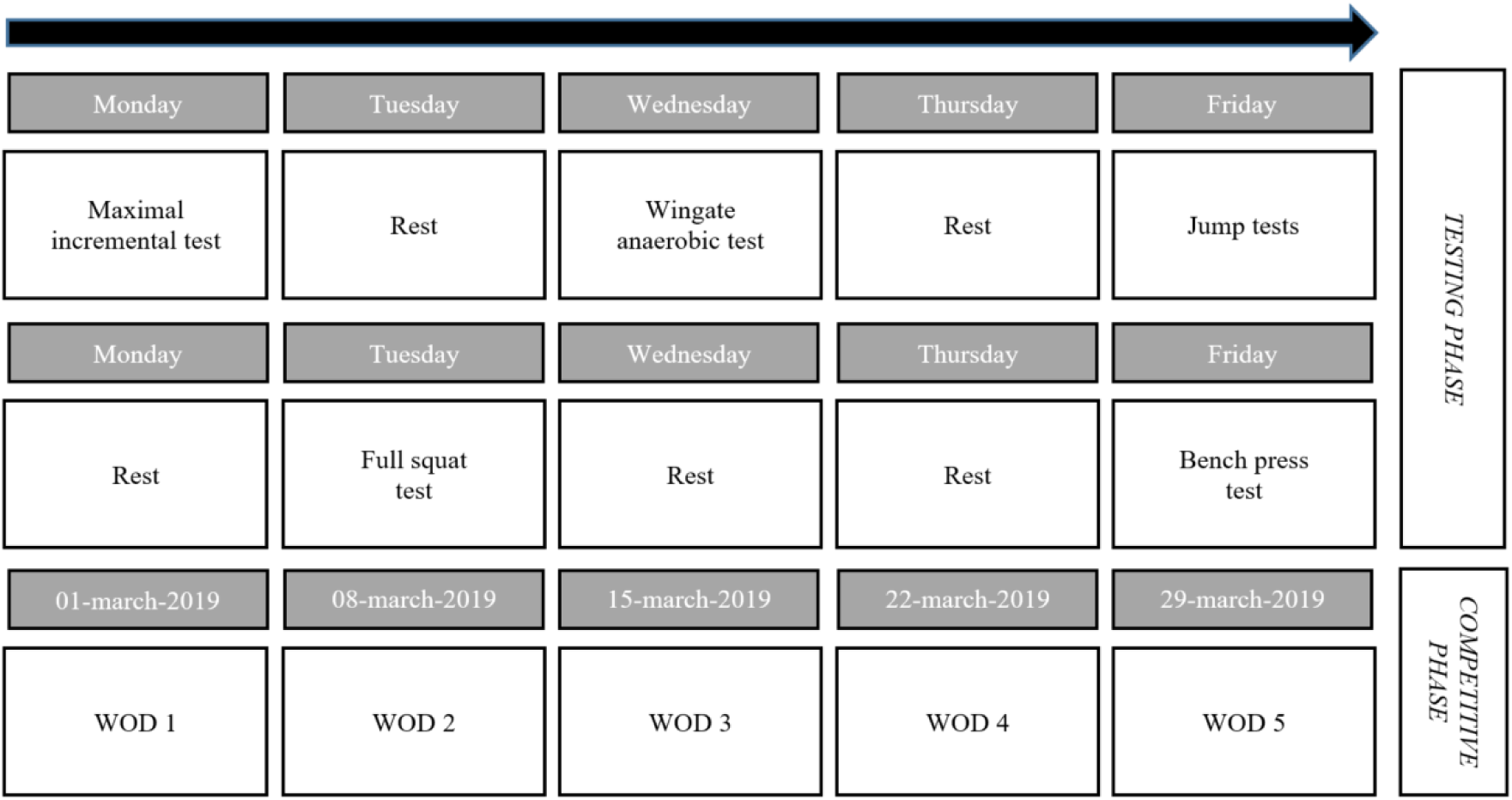
Schematic representation of the study protocol. WOD: workout of the day

### Subjects

Fifteen male athletes recruited from a local CrossFit center volunteered to participate in the study (descriptive data can be seen in Table 1). Inclusion criteria were having ≥ 1 year of experience in CrossFit, being familiar with the WAnT and the bench press and full squat exercises, and training CrossFit ≥ 3 times per week during the preceding year. During the study, participants maintained their regular training program and dietary pattern but were required to refrain from exercising at least 24 hours before each testing session or WOD, as well as from consuming ergogenic aids or stimulants (e.g., creatine, caffeine) during this period. The study was approved by the Institutional Review Board of “Hospital Universitario Fundación Alcorcón” (19/51) and all participants provided written informed consent, also were informed of the benefits and risks of the investigation prior to signing the consent.

**Table 1.** Descriptive characteristics of study participants

|  | All subjects<br>(n=15) | HP<br>(n=7) | LP<br>(n=8) | p-value | ES | Inference |
| --- | --- | --- | --- | --- | --- | --- |
| Age (years) | 35 ± 9 | 32 ± 10 | 38 ± 8 | 0.254 | 0.62 | Unclear |
| Height (cm) | 177 ± 5 | 176 ± 4 | 178 ± 6 | 0.468 | 0.41 | Unclear |
| Weight (kg) | 81 ± 9 | 78 ± 2 | 84 ± 11 | 0.161 | 0.94 | Very unlikely |
| BMI (kg·m<sup>2</sup>) | 25.9 ± 1.5 | 25.2 ± 0.6 | 26.5 ± 1.8 | 0.091 | 1.09 | Very unlikely |
| Experience (months) | 40 ± 27 | 30 ± 15 | 49 ± 33 | 0.192 | 0.78 | Very unlikely |

Data are Mean ± SD. Abbreviations: ES, effect size; HP, high-performance group; LP, low-performance group

## Measures

### Lower- and upper-body strength and power tests

Participants performed an incremental load-free test (i.e., not performed on a guided machine) for both the full squat and bench press exercises. Bar mean propulsive velocity (MPV) during the concentric phase was measured with a linear position transducer (Chronojump, Boscosystem, Spain), and power was calculated based on the total mass moved (sum of the subject’s body mass and the external load for the full squat, and only the external load for the bench press). The initial weight was 20 kg (i.e., only the bar), and the load was increased in 10–15 kg increments until a constant decrease in MPV was observed. Tests were deemed concluded when MPV decreased to 0.6 m∙s^−1^ for the full squat (6) and 0.4 m∙s^−1^ for the bench press (7). A three-minute rest was allowed between loads. Athletes performed three repetitions with each load, but only the best one (based on the mean concentric propulsive power) was used for analysis. The maximum mean concentric propulsive power (Pmax) registered during the incremental test was used for analysis as absolute (W) and relative (W∙kg^−1^) values.

The 1RM was calculated for each exercise based on the individual force-velocity relationship through linear interpolation, assuming that it was attained with an MPV of 0.30 m∙s^−1^ for the full squat (6) and 0.16 m∙s^−1^ for the bench press (7). According to recent studies, this method provides an accurate estimate of the actual 1RM (8,9). We checked that the linear regression accurately fitted the load-velocity data by examining the correlation coefficients (R^2^=0.97 ± 0.02 and 0.98 ± 0.02 for the full squat and the bench press, respectively). The 1RM was analyzed both as absolute (kg) and relative (% body weight) values.

### Jump performance

Jump performance was measured by means of an optoelectric cell system (Optojump, Microgate, Bolzano, Italy) while participants performed squat (SJ), contermovement (CMJ), and drop jumps (DJ). They performed three trials for each type of jump, and the mean of the three trials was used for analysis. Participants were instructed to place their hands on their hips while performing the jumps. During the SJ they performed a downward movement to reach 90º of knee flexion, stopped at that position for ~2 seconds, and then tried to reach the maximum jump height without performing any countermovement. During CMJ they performed the same procedure, but no stop was made at 90º of knee flexion and countermovement was allowed. For the DJ, they stepped from a 40-cm-height bench and jumped as high as possible with the minimal possible ground contact time. Reactive strength index (RSI) was calculated as jump height in the DJ divided by contact time. During all jumps participants were instructed not to flex their knees during the flight or the landing phase to avoid overestimation of flight time.

### Maximal incremental test

Participants performed an incremental running test on a treadmill (HP Cosmos Quasar, Nussdorf-Traunstein, Germany) for the determination of the first (VT1) and second (VT2) ventilatory thresholds, as well as for the determination of VO_2max_. After a 3-minute warm-up at 5 km•h^−1^, the test started at 6 km•h^−1^ and speed was increased by 0.25 km•h^−1^ every 15 seconds, keeping the inclination steady at 1% during the entire test (10). The test was terminated upon volitional fatigue or when participants could not maintain the required speed. Gas exchange data were collected continuously using a breath-by-breath system (Ultima Series Medgraphics, Cardiorespiratory Diagnostics, Saint Paul, MN). VT1, VT2 and VO_2max_ were determined as explained elsewhere (11). Peak velocity (V_peak_) was defined as the highest velocity attained during the test. We also assessed the mean muscle oxygen saturation (SmO_2_) of the right *vastus lateralis* during the incremental test by means of near infrared spectrometry (Humon, Cambridge, MA) (12).

### Wingate Anaerobic Test

Participants performed the WAnT on a cycle-ergometer (Monark, 818 E, Varberg, Sweden) as explained elsewhere (13). After a warm-up, pedaling with a resistance of ~2% of their body weight and a cadence of 70–90 rpm, they completed a 30-second all-out test with a resistance of 7.5% of their body weight (13). The mean (MPO) and peak (PPO) power output were determined as the average PO attained during the test and the highest PO achieved during 3 consecutive seconds, respectively. The fatigue slope (FS) was computed with the following equation (13):

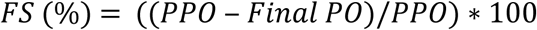

The mean SmO_2_ of the right *vastus lateralis* muscle was measured during the WAnT as mentioned for the incremental running test. Fingertip capillary blood samples (0.5 μl) were taken at baseline, and at 0, 3, 5 and 10 minutes after the test, and lactate concentration was quantified using a portable lactate analyzer (Lactate Scout, SensLab GmbH, Germany). The highest lactate value recorded for each participant was considered as the lactate peak ([La^−^]_peak_).

### CrossFit performance

The specific details of the five WODs used in this study, known as 19.1, 19.2, 19.3, 19.4 and 19.5, can be seen at https://games.crossfit.com/workouts/open/2019. They are briefly explained below:

- WOD 1. Participants had to complete in 15 minutes as many rounds as possible of 19 wall-ball shots (20-lb ball to a 10-foot target) and 19 calories of rowing.
- WOD 2. Participants had to complete in 8 minutes 25 toes-to-bar, 50 double-unders, 15 squat cleans (135 lb), 25 toes-to-bar, 50 double-unders, and 13 squat cleans (185 lb). If they completed these exercises before 8 minutes, 4 further minutes were added and they had to perform 25 toes-to-bar, 50 double-unders, and 11 squat cleans (225 lb). If they completed again these exercises before 12 minutes, 4 minutes were added and they had to perform 25 toes-to-bar, 50 double-unders, and 9 squat cleans (275 lb). If completed before 16 minutes, 4 additional minutes were added and they performed 25 toes-to-bar, 50 double-unders, and 7 squat cleans (315 lb). The maximum time allowed was 20 minutes.
- WOD 3. Participants had a maximum of 10 minutes to complete 200-foot dumbbell (50 lb) overhead lunges, 50 dumbbell (50 lb) box step-ups (24-inch box), 50 strict handstand push-ups, and 200-foot handstand walk in the minimum time possible.
- WOD 4. Participants had a maximum of 12 minutes to complete 3-rounds of 10 snatches (95 lb) and 12 bar-facing burpees in the minimum time possible. They then rested for 3 minutes and continued with 3 rounds of 10 bar muscle-ups and 12 bar-facing burpees, which they had to complete in the minimum time possible.
- WOD 5. Participants had 20 minutes to perform 33 thrusters (95 lb) and 33 chest-to-bar pull-ups, followed by 27, 21, 15, and 9 reps of the same sequence (i.e., same number for thrusters and chest-to-bar pull-ups).

Participants were then ranked in positions (i.e., from 1 to 15) depending on their performance (time or repetitions, depending on the WOD) in each WOD. They also received a score after each WOD depending on their classification within the group (one point for the first position, two points for the second one, and so on), and were then ranked for overall CrossFit performance considering the sum of the scores attained in the five WODs (with lower scores reflecting a better performance). Only those participants who performed all the WODs were included in the analyses. Participants were divided by the median into a low-performance (LP) and a high-performance (HP) group attending to the final ranking.

### Statistical analysis

Normal distribution (Shapiro-Wilk test) and homoscedasticity (Levene’s test) of the data were checked before any statistical treatment. Spearman’s rank correlation coefficients (ρ) were calculated to analyze the relationship between each variable and the position (i.e., ranking) within the group. Least-squares multiple linear regression analysis was used to analyze which variables from those that appeared significant in simple linear regression could together predict overall CrossFit performance. Correlation coefficients of 0.1, 0.3, 0.5, 0.7 and 0.9 were considered small, moderate, large, very large and extremely large, respectively (14). Independent samples t-tests were conducted to analyze differences between HP and LP groups. The likelihood of finding differences were assessed using magnitude-based inferences and considered as follows: <0.5%, most unlikely; 0.5–5%, very unlikely; 5–25%, unlikely; 25–75%, possibly; 75–95%, likely; 95–99.5%, very likely; >99.5, most likely (14). If the chances of having better and poorer results were both >5%, the difference was considered unclear. The magnitude of the differences between groups was assessed through the computation of effect sizes (ES; Cohen’s d), which were considered trivial (<0.2), small (<0.6), moderate (<1.2) large (<2.0) or very large (<4.0) (14). Analyses were conducted with a statistical software package (SPSS 23.0, IBM; Armonk, NY) and an Excel Spreadsheet (15), setting an α level of p<0.05 as the minimal level of significance.

## Results

Differences between the HP and LP groups are shown in Table 1 and Table 2. No significant differences were observed for age, anthropometrical variables or training experience (p>0.05) (Table 1).

**Table 2.** Differences between high- and low-performance groups

| Test | Variable | HP (n=7) | LP (n=8) | p-value | ES | Inference |
| --- | --- | --- | --- | --- | --- | --- |
| JUMP TEST | SJ (cm) | 39 ± 5 | 32 ± 3 | **0.006\*** | 1.73 | Very likely |
|  | CMJ (cm) | 42 ± 5 | 37 ± 6 | 0.105 | 0.91 | Very likely |
|  | RSI (cm/ms) | 0.92 ± 0.22 | 0.73 ± 0.11 | 0.052 | 1.15 | Likely |
| WINGATE TEST | MPO (W) | 671 ± 39 | 639 ± 70 | 0.306 | 0.59 | Unclear |
|  | PPO (W) | 823 ± 58 | 788 ± 86 | 0.367 | 0.50 | Unclear |
|  | MPO (W·kg<sup>-1</sup>) | 8.6 ± 0.6 | 7.6 ± 0.8 | **0.019\*** | 1.43 | Very likely |
|  | PPO (W·kg<sup>-1</sup>) | 10.6 ± 0.7 | 9.4 ± 1.1 | **0.030\*** | 1.31 | Very likely |
|  | FS (%) | 37 ± 11 | 35 ± 8 | 0.612 | 0.27 | Unclear |
|  | SmO<sub>2</sub> (%) | 67 ± 7 | 69 ± 6 | 0.601 | 0.29 | Unclear |
|  | [La<sup>-</sup>]<sub>peak</sub> (mmol·l<sup>-1</sup>) | 15.3 ± 2.8 | 13.4 ± 4.3 | 0.325 | 0.32 | Unclear |
| INCREMENTAL TEST | VO<sub>2max</sub> (ml·kg<sup>-1</sup>·min<sup>-1</sup>) | 55 ± 5 | 49 ± 4 | **0.022\*** | 1.35 | Very likely |
|  | Vpeak (km·h<sup>-1</sup>) | 16.1 ± 1.0 | 14.7 ± 0.9 | **0.011\*** | 1.52 | Very likely |
|  | VT1 (km·h<sup>-1</sup>) | 10.1 ± 0.6 | 9.9 ± 0.9 | 0.619 | 0.27 | Unclear |
|  | VT2 (km·h<sup>-1</sup>) | 15.1 ± 1.1 | 14.2 ± 1.1 | 0.128 | 0.84 | Likely |
|  | SmO<sub>2</sub> (%) | 64 ± 8 | 66 ± 10 | 0.733 | 0.19 | Unclear |
| STRENGTH TEST | 1RM BP (kg) | 105 ± 11 | 95 ± 21 | 0.279 | 0.63 | Unclear |
|  | 1RM BP (%BW) | 1.35 ± 0.13 | 1.13 ± 0.22 | **0.035\*** | 1.27 | Very likely |
|  | Pmax BP (W) | 554 ± 104 | 505 ± 140 | 0.463 | 0.40 | Unclear |
|  | Pmaxrel BP (W·kg<sup>-1</sup>) | 7.12 ± 1.40 | 5.96 ± 1.30 | 0.121 | 0.86 | Likely |
|  | 1RM FS (kg) | 133 ± 19 | 128 ± 28 | 0.704 | 0.21 | Unclear |
|  | 1RM FS (%BW) | 1.71 ± 0.29 | 1.52 ± 0.22 | 0.164 | 0.76 | Likely |
|  | Pmax FS (W) | 1436 ± 300 | 1553 ± 529 | 0.616 | 0.28 | Unclear |
|  | Pmaxrel FS (W·kg<sup>-1</sup>) | 13.5 ± 1.7 | 13.5 ± 1.7 | 0.916 | 0.06 | Unclear |

Data are Mean ± SD. Abbreviations: HP, high-performance group; LP, low-performance group; CMJ, contramovement jump; FS, fatigue slope; [La<sup>-</sup>]<sub>peak</sub>, lactate peak; MPO, mean power output; PPO, peak power output; RSI, reactive strength index; SJ, squat jump; SmO<sub>2</sub>, muscle oxygen saturation; Pmax BP, maximum power bench press; Pmax FS, maximum power full squat; 1RM FS, 1-Repetition maximum full squat; 1RM BP, 1-Repetition maximum bench press; VO<sub>2max</sub>, maximum oxygen consumption; Vpeak, peak speed; VT1, speed at ventilatory threshold 1; VT2, speed at ventilatory threshold 2; Significant differences between groups are highlighted in bold.

We found that jump performance (i.e., JS, CMJ and RSI) was likely or very likely to be higher in the HP group than in the LP group, although these differences only reached statistical significance for the JS test (Table 2). The HP group also presented with a higher PPO and MPO during the WAnT, but only when expressed in relative values (Table 2), as well as with a higher VO_2max_ and Vpeak (Table 2). No differences (p>0.05) were observed, however, for VT1 or VT2. Lastly, the HP group showed a higher relative 1RM in the bench press exercise, but no between-group differences were observed for absolute 1RM or for any of the analyzed variables in the full squat (Table 2).

The relationships between the variables registered in the different tests, and the performance in each WOD or the final position within the group, are shown in Table 3. Jump ability was the best predictor of CrossFit performance, with all jump-related variables (i.e., SJ, CMJ and RSI) largely associated with ≥ 4 out of the 5 WODs performed, as well as to the final ranking (Table 3). The same trend was observed for the relative – but not absolute – PPO and MPO during the WAnT, which were also largely related to performance in ≥ 4 WODs as well as to the final ranking (Table 3). VO_2max_ and Vpeak were also significantly and strongly related to performance in 3 WODs and to the final ranking (Table 3). Lastly, relative – but not absolute – upper and lower-body maximal strength (i.e., 1RM for the bench press and the full squat, respectively, analyzed as a % body weight) were largely related to performance in ≥ 3 WODs as well as to the final ranking (Table 3). A significant relationship was also observed for the maximum power in the bench press exercise, but this relationship remained significant in only one of the five analyzed WODs.

**Table 3.** Relationship between the different physiological variables and CrossFit performance

| Test | Variable | WOD 1 | WOD 2 | WOD 3 | WOD 4 | WOD 5 | Final score |
| --- | --- | --- | --- | --- | --- | --- | --- |
|  |  | r | r | r | r | r | r |
| JUMP TEST | SJ (cm) | -.32 | **-.69\*\*** | **-.73\*\*** | **-.69\*\*** | **-.73\*\*** | **-.74\*\*** |
|  | CMJ (cm) | **-.56\*** | **-.55\*** | **-.51\*** | **-.55\*** | **-.65\*\*** | **-.62\*\*** |
|  | RSI (cm/ms) | **-.59\*** | **-.89\*\*** | **-.70\*\*** | **-.72\*\*** | **-.62\*** | **-.74\*\*** |
| WINGATE TEST | MPO (W) | **-.54\*** | -.41 | -.25 | -.32 | -.32 | -.43 |
|  | PPO (W) | -.49 | -.35 | -.16 | -.26 | -.09 | -.28 |
|  | FS (%) | -.06 | .70 | .01 | -.07 | .13 | .11 |
|  | MPO (W·kg<sup>-1</sup>) | -.48 | **-.74\*\*** | **-.63\*\*** | **-.69\*\*** | **-.74\*\*** | **-.75\*\*** |
|  | PPO (W·kg<sup>-1</sup>) | **-.63\*** | **-.62\*** | -.52 | **-.64\*** | **-.59\*** | **-.61\*** |
|  | SmO<sub>2</sub> (%) | -.24 | .30 | -.06 | -.20 | -.00 | .16 |
|  | [La<sup>-</sup>]<sub>peak</sub> (mmol·l<sup>-1</sup>) | -.12 | **-.53\*** | -.34 | -.44 | -.38 | -.40 |
| INCREMENTAL TEST | VO<sub>2max</sub> (ml·kg<sup>-1</sup>·min<sup>-1</sup>) | **-.60\*** | -.47 | -.51 | **-.55\*** | **-.63\*** | **-.58\*** |
|  | Vpeak (km·h<sup>-1</sup>) | -.34 | -.45 | **-.55\*** | **-.57\*** | **-.71\*\*** | **-.68\*** |
|  | VT1(km·h<sup>-1</sup>) | -.00 | .09 | .10 | .03 | -.23 | -.09 |
|  | VT2 (km·h<sup>-1</sup>) | -.25 | -.38 | -.39 | -.34 | -.28 | -.38 |
|  | SmO<sub>2</sub> (%) | -.21 | -.21 | -.04 | -.13 | -.02 | -.01 |
| STRENGTH TEST | 1RM BP (kg) | -.20 | -.15 | -.27 | -.19 | -.13 | -.34 |
|  | 1RM BP (%BW) | -.28 | -.50 | **-.71\*\*** | **-.65\*** | **-.65\*\*** | **-.60\*\*** |
|  | Pmax BP (W) | -.45 | -.22 | -.15 | -.15 | -.22 | -.36 |
|  | Pmax BP (W·kg<sup>-1</sup>) | -.50 | -.44 | -.42 | -.41 | **-.53\*** | **-.59\*** |
|  | 1RM FS (kg) | -.38 | -.39 | -.38 | -.41 | **-.57\*** | -.35 |
|  | 1RM FS (%BW) | -.41 | **-.56\*** | **-.54\*** | **-.60\*** | **-.78\*\*** | **-.66\*\*** |
|  | Pmax FS (W) | -.35 | -.25 | -.02 | -.01 | .00 | -.05 |
|  | Pmax FS (W·kg<sup>-1</sup>) | -.34 | -.34 | -.12 | -.12 | -.17 | -.29 |

Abbreviations: CMJ, contramovement jump; FS, fatigue slope; [La-]peak, lactate peak; MPO, mean power output; PPO, peak power output; RSI, reactive strength index; SJ, squat jump; SmO<sub>2</sub>, muscle oxygen saturation; Pmax BP, maximum power bench press; Pmax FS, maximum power full squat; 1RM FS, 1-Repetition maximum full squat; 1RM BP, 1-Repetition maximum bench press; VO<sub>2max</sub>, maximum oxygen consumption; Vpeak, peak speed; VT1, speed at ventilatory threshold 1; VT2, speed at ventilatory threshold 2; Performance was considered as the position (1–15) attained within the group in each workout of the day (WOD), and the final score is the sum of the points achieved in each WOD for each athlete. A higher position indicates a worse performance. Significant correlations are highlighted in bold: \*p<0.05, \*\*p<0.01.

Multiple linear regression analysis showed that the combination of VO_2max_, SJ, and RSI accounted for 81% of the performance variance (R_2_=80.7 %, p=0.0003). The following equation determined the relationship:

CrossFit performance (overall score) = 192.682 – 38.3259*RSI (cm/ms) – 1.51687*SJ (cm) – 1.31099*VO_2max_ (ml∙kg^−1^∙min^−1^)

## Discussion

The aim of the present study was to analyze the relationship between CrossFit performance and a variety of physiological markers related to ‘aerobic’ and ‘anaerobic’ capacity, strength, and power. Our results show that CrossFit performance is dependent on a range of physiological ‘domains’, with power-(e.g., jump ability and power during the WAnT), strength-(e.g., relative 1RM) and aerobic-related markers (e.g., VO_2max_ and Vpeak) all being related to overall CrossFit Performance. Indeed, these variables – particularly jump performance and relative power during the WAnT – appeared as strong predictors of CrossFit performance in most of the WODs performed (~3–4 out of 5), and the combination of power-(i.e., SJ and RSI) and aerobic-(VO_2max_) related markers together explained most of the variance in overall CrossFit performance (81%).

Research in CrossFit performance has exponentially grown in recent years, but there is still scarce information on the performance determinants of this sport (1). In the present study lower-body muscle – as measured by jump ability and PPO during the WAnT – appeared as one of the strongest predictors of performance. Other studies have previously found a relationship between lower-body muscle power and CrossFit performance. For instance, we recently observed that power-related indices measured on the squat test were related to a greater performance in most of the WODs analyzed (3). Similarly, in agreement with our findings, other authors recently reported that the PPO measured during a WAnT was related to performance in two out of the four WODs analyzed (5). Of note, some CrossFit exercises included in this study such as singles or double unders involve repeated jumps. Moreover, jump ability has been related to performance in other exercises such as the loaded squat jump (16,17). On the other hand, the PPO measured during the WAnT is also related to key athletic actions including jumping or sprinting (18), which are usually present in CrossFit WODs. Thus, the assessment of lower-body muscle power can provide valuable information on CrossFit athletic performance. If must be noted that, contrary to a recent study (3), we did not observe a clear relationship between power indices measured during the full squat and CrossFit performance, which can be potentially due to the lower velocity attained during this exercise compared with other power actions such as the WAnT or unloaded jumps. Future research should confirm the validity of power measures obtained during the full squat for the prediction of CrossFit performance.

The relative – but not absolute - maximum strength of both the upper and lower limbs was related to CrossFit performance, which reflects the importance of body weight in most exercises (e.g. wall-ball shots, squat cleans, overhead lunges, handstand walk).

In most CrossFit exercises, athletes not only have to lift or throw an external load, but also their own body mass. For this reason, as in other sports, trying to reach a balance between maximum strength and body mass will be of paramount importance (19), although in this case the importance can likely differ depending on the WOD performed (2,3,5).

Finally, an interesting finding of this study was that both ‘aerobic-’ and ‘anaerobic’-related markers were related to CrossFit performance, as measured by the VO_2max_ and Vpeak during the incremental test, and the MPO or PPO during the WAnT. These results are in agreement with those of previous studies, in which VO_2max_ was related to performance in CrossFit WODs such as Fran, Cindy or Grace (5). Moreover, other authors have also found a relationship between performance on the WAnT and performance in different types of WODs (20).

Some limitations of this study must be acknowledged, such as its cross-sectional nature, which precludes us from knowing whether enhancing any of the analyzed variables would result in an improved CrossFit performance. Moreover, the characteristics and exercises of the CrossFit Open Games change each year. Thus, the present findings might not be necessarily applicable to WODs other than those analyzed here. It must be noted, however, that some of the analyzed markers were related to performance in WODs that included a great variety of exercises (e.g., wall-ball shots, burpees, push ups, or lunges), which suggest that this association might also be observed in other WODs.

## Conclusions

In summary, the present study shows that CrossFit performance is at least partly dependent of a variety of power-(e.g., jump ability and power during the WAnT), strength-(e.g., relative 1RM for full squat and bench press) and aerobic-related (e.g., VO_2max_ and Vpeak) markers, which reflects the complexity of this sport. Measures of lower-body muscle power (particularly jump ability) and aerobic capacity (as measured through the VO_2max_) together explained most of the variance in overall CrossFit performance, and thus these tests could be potentially used for the indirect assessment of CrossFit performance.

## Author contributions

**Rafael Martínez-Gómez**: conceptualization, data curation, investigation, methodology, project administration, resources, writing.

**Pedro L. Valenzuela**: conceptualization, data curation, investigation, methodology, project administration, resources, writing.

**Lidia B. Alejo**: conceptualization, data curation, investigation, methodology, project administration, resources, writing.

**Jaime Gil-Cabrera**: data curation, investigation, resources, validation. **Almudena Montalvo-Pérez**: data curation, investigation, resources, validation. **Eduardo Talavera**: data curation, investigation, resources, validation.

**Alejandro Lucia**: data curation, investigation, resources, validation.

**Susana Moral-González**: conceptualization, data curation, investigation, resources, validation.

**David Barranco-Gil**: conceptualization, data curation, formal analysis, methodology, investigation, resources, supervision, validation.

## Acknowledgments

We also thank all the participants as well as CrossFit Las Rozas (Madrid, Spain) and the head coach Manuel Gómez Martin. The work of Pedro L. Valenzuela is supported by University of Alcalá (FPI2016). Research by Alejandro Lucia is funded by the Spanish Ministry of Economy and Competitiveness and Fondos Feder [grant #PI PI18/00139].

## References

1. Claudino J, Gabbett T, Bourgeois F, Souza H, Miranda R, Mezêncio B, et al. CrossFit Overview: Systematic Review and Meta-analysis. Sports Med - Open 2018 Dec;4(1):1–14.

2. Butcher SJ, Neyedly TJ, Horvey KJ, Benko CR. Do physiological measures predict selected CrossFit(®) benchmark performance? J Sports Med 2015;6:241–247.

3. Martínez-Gómez R, Valenzuela PL, Barranco-Gil D, Moral-González S, García-González A, Lucia A. Full-Squat as a Determinant of Performance in CrossFit. Int J Sports Med 2019 Jul 10.

4. Bellar D, Hatchett A, Judge LW, Breaux ME, Marcus L. The relationship of aerobic capacity, anaerobic peak power and experience to performance in CrossFit exercise. Biol. Sport 2015 Nov;32(4):315–320.

5. Dexheimer JD, Schroeder ET, Sawyer BJ, Pettitt RW, Aguinaldo AL, Torrence WA. Physiological Performance Measures as Indicators of CrossFit ® Performance. Sports 2019 Apr 22;7(4):93.

6. Conceição F, Conceição F, Fernandes J, Lewis M, Gonzaléz-Badillo JJ, Jimenéz-Reyes P. Movement velocity as a measure of exercise intensity in three lower limb exercises. Journal of Sports Sciences 2016 Jun 17;34(12):1099–1106.

7. J.J. González-Badillo LS. Movement Velocity as a Measure of Loading Intensity in Resistance Training. Int J Sports Med 2010 February 23;31:347–352.

8. Garcia-Ramos A, Jaric S. Feasibility of the Two-Point Method for Determining the One-Repetition Maximun in the Bench Press Exercise. Strength Cond J 2018 Apr 1;40(2):54–66.

9. García-Ramos A, Pestaña-Melero FL, Pérez-Castilla A, Rojas FJ, Haff GG. Differences in the Load–Velocity Profile Between 4 Bench-Press Variants. International journal of sports physiology and performance 2018 Mar 1;13(3):326–331.

10. Andrew M. Jones and Jonathan H. Doust. A 1% treadmill grade most accurately reflects the energetic cost of outdoor running. J Sports Sci 1996;14:321–327.

11. Lucía A, Pardo J, Durántez A, Hoyos J, Chicharro JL. Physiological differences between professional and elite road cyclists. Int J Sports Med 1998 Jul;19(5):342–348.

12. Farzam P, Starkweather Z, Franceschini MA. Validation of a novel wearable, wireless technology to estimate oxygen levels and lactate threshold power in the exercising muscle. Physiol Rep 2018 Apr;6(7):e13664.

13. Bar-Or O. The Wingate anaerobic test. An update on methodology, reliability and validity. Sports Med 1987 Nov;4(6):381–394.

14. Hopkins WG, Marshall SW, Batterham AM, Hanin J. Progressive Statistics for Studies in Sports Medicine and Exercise Science. Medicine and science in sports and exercise 2009 Jan;41(1):3–13.

15. Hopkins W. A spreadhseet to compare means of two groups. SportsSci 2007;11:22–23.

16. Loturco I, Pereira LA, Moraes JE, Kitamura K, Cal Abad CC, Kobal R, et al. Jump-Squat and Half-Squat Exercises: Selective Influences on Speed-Power Performance of Elite Rugby Sevens Players. PloS one 2017;12(1):e0170627.

17. Comfort P, Stewart A, Bloom L, Clarkson B. Relationships Between Strength, Sprint, and Jump Performance in Well-Trained Youth Soccer Players. J Strength Cond Res 2014 Jan;28(1):173–177.

18. Hoffman JR, Epstein S, Einbinder M, Weinstein Y. A Comparison Between the Wingate Anaerobic Power Test to Both Vertical Jump and Line Drill Tests in Basketball Players. J Strength Cond Res 2000 Aug;14(3):261–264.

19. Andersen E, Lockie RG, Dawes JJ. Relationship of Absolute and Relative Lower-Body Strength to Predictors of Athletic Performance in Collegiate Women Soccer Players. Sports (Basel, Switzerland) 2018 Sep 29;6(4):106.

20. José Luis Maté-Muñoz, Juan H Lougedo, Manuel Barba, Pablo García-Fernández, Manuel V Garnacho-Castaño, Raúl Domínguez. Muscular fatigue in response to different modalities of CrossFit sessions. PLoS One 2017 Jul 1;12(7):e0181855.

